# Deep Visual Proteomics Unveils Precision Medicine Insights in Composite Small Lymphocytic and Classical Hodgkin Lymphoma

**DOI:** 10.1101/2024.06.12.598635

**Authors:** Xiang Zheng, Lars Møller Pedersen, Michael Bzorek, Andreas Mund, Lise Mette Rahbek Gjerdrum, Matthias Mann

## Abstract

Coexistence of two cancer types in the same organ presents challenges for clinical decision-making, calling for personalized treatment strategies. Deep Visual Proteomics (DVP) combines AI driven single cell type analysis with laser microdissection and ultrasensitive mass spectrometry. In a composite case of classical Hodgkin lymphoma (cHL) and small lymphocytic lymphoma (SLL) in a single patient, we investigated the potential of DVP to inform precision oncology. We quantified the proteomic landscapes in the cHL and SLL to a depth of thousands of proteins. Our analysis revealed distinct proteome profiles in cHL and SLL populations, highlighting their clonal unrelatedness. Our data suggested standardized chemotherapy and interleukin-4 inhibition as potential strategies to manage chemo-resistance – instead of bone marrow transplantation. DVP highlighted minichromosome maintenance protein and proteasome inhibitors for cHL and H3K27 methylation and receptor tyrosine kinase inhibitors for SLL as subtype-specific treatments. Thus cell-type specific insights of DVP can guide personalized oncological treatments.

## Introduction

Mass spectrometry (MS)-based proteomics has made great advances in recent years and is an increasingly powerful complement to the spatial omics toolbox ^1^. While previously being restricted to bulk tissue analysis, technological advances have now brought the capability to analyze cellular heterogeneity across tissues and thereby the global determination of molecular features of individual cell types ^2^. In particular, our group has recently described Deep Visual Proteomics (DVP), a novel technology for spatial single-cell-type proteome analysis, which combines high-resolution imaging, AI-based single-cell phenotyping, automated single-cell laser microdissection and ultrasensitive mass spectrometry (MS) ^3^. Given DVP’s capacity to provide a detailed proteomic view of individual cell types, it would be interesting to evaluate its therapeutic relevance in personalized treatment options and disease management.

Composite lymphoma provides an ideal context for exploring the potential of DVP in advancing precision medicine strategies. The co-occurrence of classical Hodgkin lymphoma (cHL) and small lymphocytic lymphoma (SLL) or chronic lymphocytic leukemia (CLL) is a rare phenomenon (with incidence less than 1% of all newly diagnosed lymphomas per year), however, treatment approaches for composite lymphomas remain elusive ^4, 5^. Typically, more than 90% of lymphoproliferative malignancies arise from B cells, while the remainder originates from T cells or NK cells ^6^. Composite lymphomas may involve classical Hodgkin lymphoma (cHL) and a non-Hodgkin lymphoma (NHL) within the same patient ^7^. The development of composite cHL and NHL typically stems from the evolution of two distinct malignant germinal center B cell clones, originating from a shared premalignant precursor cell with common mutations. Subsequently, these daughter cells acquire additional unique mutations, leading to the emergence of two separate B-cell neoplasms ^7, 8^. The occurrence of aggressive cHL in individuals previously diagnosed with chronic lymphocytic leukemia (CLL) or SLL, known as the cHL variant of Richter’s transformation, is exceptionally rare ^9, 10^. The median time from CLL/SLL diagnosis to diagnosis of HL has been reported as 6 years, and the majority of patients have received treatment prior to the time of transformation ^11^. Only a small subset of patients has HL and CLL/SLL diagnosed simultaneously. Two-year overall survival has been estimated at 57%, with a subgroup that tolerates curative-intent therapy having a prognosis similar to patients with *de novo* HL without transformation ^11, 12^. In general, patients with HL transformation have a more favorable prognosis than patients with Richter transformation ^12, 13^.

Current treatment decisions for composite lymphomas primarily target the most aggressive histological subtype and usually involve a combination of chemotherapy, radiation therapy, or targeted therapies and in some cases, allogeneic bone marrow transplantation ^4, 9, 10^. Given the diverse clinical behaviors, poor prognosis, comorbidities and patient fitness of composite lymphomas, personalized treatment strategies are clearly needed to enhance clinical outcomes. In this study, we employ DVP to investigate a unique case involving concurrent cHL and SLL, identified at the time of primary diagnosis, to explore the capacity of DVP to provide insights into precision medicine strategies.

## Results

### Pathology workup as the initiation point for the DVP workflow

In line with the DVP workflow ^3^, we initiated our analysis by identifying and categorizing individual cell types using immunohistochemical staining and high-resolution imaging (**Methods**, **Figure 1A-B**). Our study focused on a composite lymphoma case with cervical lymphadenopathy of a 71 year old female with no previous treatment. Analysis of the excised lymph node revealed the concurrent cHL and SLL using multiple immunohistochemical markers. The cHL exhibited characteristic Hodgkin and Reed-Sternberg cells (HRS) with a classical immunohistochemical phenotype and a T-cell background with eosinophils, whereas the SLL component displayed small lymphocytes with clumped chromatin and a relevant immunophenotype (**Figure 1C-E; Supplementary figure 1A-B**) ^6, 14^. The cHL component exhibited a classical immunophenotype of CD30^+^CD15^+^IMP3^+^P53^+^PAX5^dim^CD20^-^. Conversely, the SLL component exhibited characteristics of CD20^+^PAX5^+^P53^dim^CD15^-^IMP3^-^ CD30^-^ small lymphocytes (**Figure 1C-E; Supplementary figure 1A-B**). An expert hematopathologist guided annotations of non-lesional areas containing germinal centers and lesional regions characterized by proliferation centers and diffuse areas containing lymphoma cells in SLL(**Figure 1C**).

**Figure 1.**
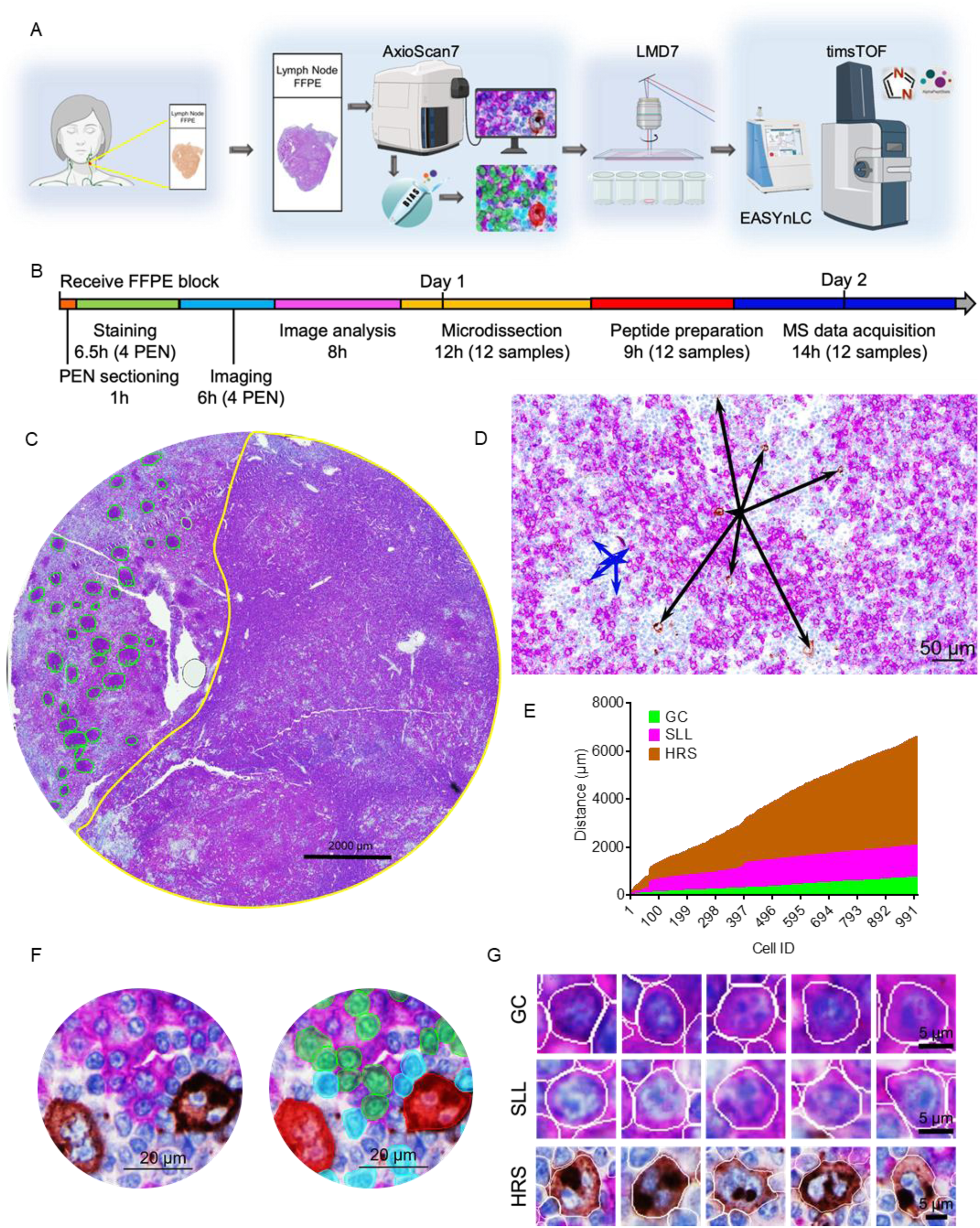
Deep Visual Proteomics workflow for a case with composite small lymphocytic lymphoma (SLL) and classical Hodgkin lymphoma (cHL). (A) Formalin fixed paraffin embedded (FFPE) tissues on membrane slides were stained using combined immunohistochemistry for CD20 and CD30, followed by high-resolution imaging, image analysis, single-cell laser microdissection and proteomics analysis (see also **Methods**). (B) The timeline shows the total duration and hands-on time of each step for the entire data acquisition. (C) Immunohistochemistry staining to identify CD20 and CD30 positive cells. The non-lesional region, containing residual germinal centers, appears in green, while the tumor region containing HRS and SLL cells is in yellow. (D) A representative image of proximity distance. Black arrows indicate the six nearest HRS to a given HRS, while blue arrows indicate the six nearest SLL to a given SLL. (E) The distance from a given GC/SLL/HRS to the nearest 1000 GC/SLL/HRS, respectively. (F) Magnified images show CD20 staining in magenta and CD30 staining in brown. A phenotype map illustrates CD20^+^CD30^-^ SLL cells with green masks, CD30^+^CD20^-^ HRS cells with red masks, and other classes with cyan masks. (G) Representative images of classification and masks for laser microdissection. CD20 staining in magenta and CD30 staining in brown. Source data are provided as a Source Data file. Abbreviations: GC, germinal center B cells; HRS: Hodgkin Reed-Sternberg cells; SLL: small lymphocytic lymphoma cells; LMD: laser micro dissection.

To facilitate single-cell microdissection, we identified B cells in residual germinal centers (GC cells) as CD20^+^CD30^-^ cells. Within the tumor region, SLL cells were CD20^+^CD30^-^ and HRS cells were CD30^+^CD20^-^ with further characteristic morphology (**Figure 1C**). The sparsity of HRS posed a challenge in accurately identifying and locating them within the tissue (**Figure 1D**). This is reflected in the fact that the distance between HRS was over twice as large as the distance between SLL (calculated from 1000 nearest HRS or SLL cells) (**Figure 1D-E**). Therefore, precise cell type identification was crucial for our analysis. To accurately identify and classify GC, SLL, and HRS cells, we trained a deep neural network based on CD20 and CD30 expression and morphological features in the BIAS software as described before ^3^ (**Figure 1F-G**). This used 10-fold cross validation, a resampling procedure, to evaluate the performance of the trained models. The accuracy of the models for predicting HRS and SLL was 98% (**Figure 1F-G**). The tumor microenvironment also comprised other cell types, including T cells (**Figure 1F; Supplementary figure 1B**). Our robust cell type identification enabled us to confidently excise single-cell-types using laser microdissection (LMD) (The average of 300 HRS shapes and 1000 SLL shapes, corresponding to 70,000 μm² surface areas, and 100 intact HRS and 600 intact SLL, respectively (**Methods**, **Figure 1G; Supplementary figure 2A-B**).

### Substantial proteome differences between HRS and SLL cells

Next, we performed single-cell laser microdissection followed by ultra-sensitive proteomics analysis to profile the two distinct cell types. To ensure reliable and accurate identification and quantification of proteins in data independent acquisition (DIA), we first generated a deep project-specific library. This contained 107,403 peptides from 9,472 protein groups, generating from data-dependent acquisition (DDA) runs of pre-fractionated samples specific to the experiment. We identified 2831, 3183 and 3090 protein groups in GC, HRS and SLL populations, respectively (**Figure 2A-B, Supplementary figure 2C-D**). Comparing HRS or SLL to GC, the top ten significantly differentially expressed proteins, nine of them have previously been linked to lymphoma: TUBB2A, KPNA2, CDK1, IRF4, FSCN1, DDX21, TSR1, collagen and C1QA (**Figure 2A-B**) ^15, 16, 17, 18, 19, 20, 21, 22, 23^. Furthermore, upregulation of SLC3A2 in SLL was in alignment with data from the Human Protein Atlas (HPA) database ^24^ (**Supplementary figure 3A**). These observations suggest that the composite lymphoma shares molecular features with cHL and NHL.

**Figure 2.**
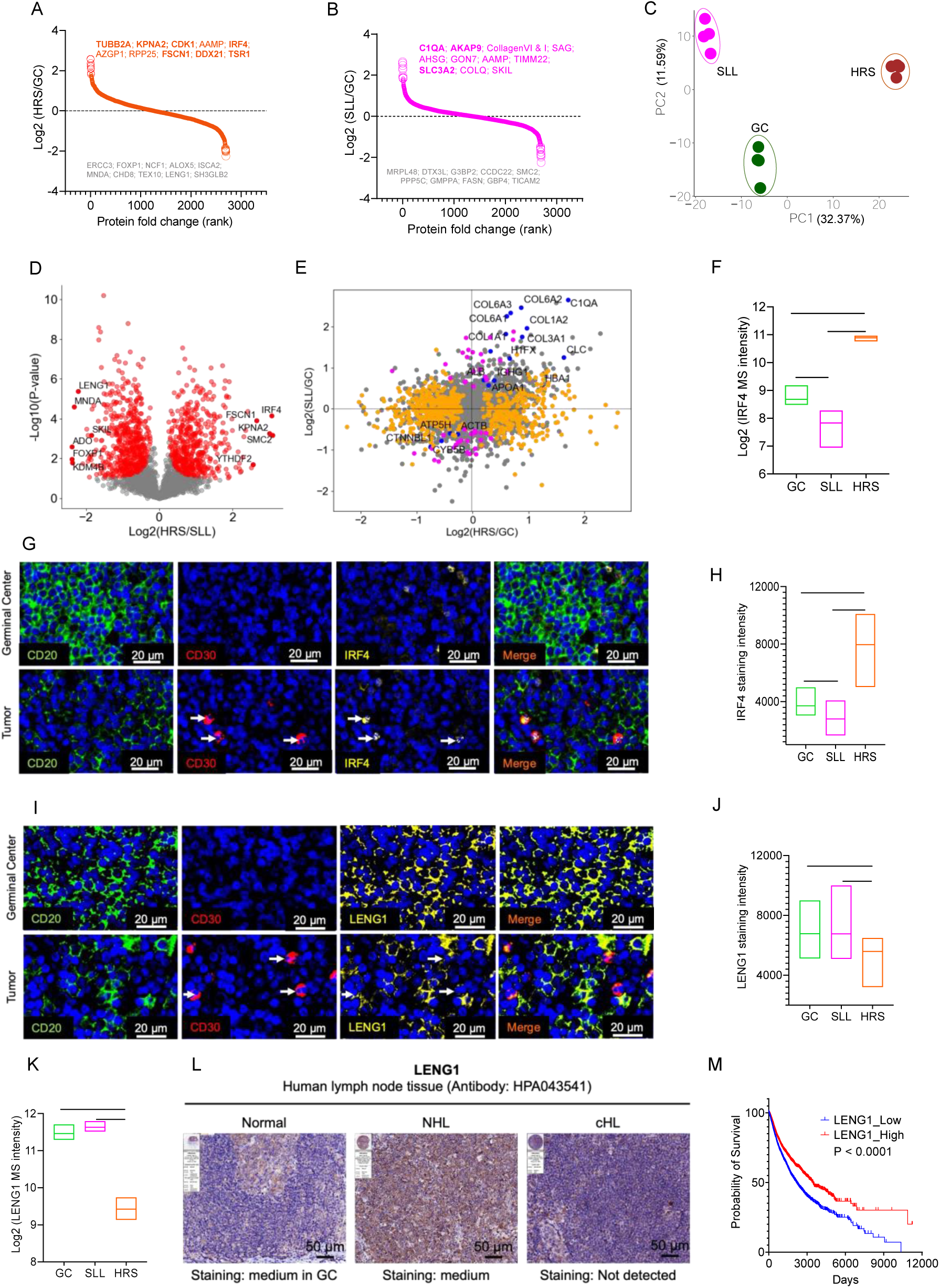
Proteomic profiling of HRS and SLL cell populations. (A) Proteins ranked by their expression differences in HRS compared to GC. The top ten differentially expressed proteins in HRS are listed, with known lymphoma-related markers in bold. (B) Same as A but for SLL compared to GC. (C) Principal component analysis of GC/HRS/SLL cell proteomes. (D) Volcano plot of the proteomic comparison between HRS and SLL. Red dots represent significantly differentially expressed proteins. Proteins with largest fold-change are annotated with their names. Statistical significance was determined using two-sided *t*-test (FDR<0.05, S0=0.1). (E) The scatter plot of the differential enrichment of proteins in the HRS and SLL populations in comparison to GC. Proteins that show significant expression in HRS (orange), SLL (magenta), and both populations (blue) are highlighted. (F) Expression of IRF4 determined by MS analysis. The box plot displays the distribution of the data from the minimum to the maximum intensity. The horizontal line within each box represents the mean value of the intensity. Statistical significance was determined using one-way ANOVA. F-value (2, 9) = 92.55; **** P < 0.0001. * P < 0.05. (G) Representative images illustrating immunofluorescence staining of CD20, CD30 and IRF4 in non-lesional germinal center regions and tumor areas. Arrows indicate HRS cells. (H) Quantification of staining intensity of IRF4 in GC, HRS and SLL (similar as in F). The horizontal line within each box represents the median value of the intensity. Statistical significance was determined using one-way ANOVA. F (2, 41786) = 4132, **** P < 0.0001. (I) Representative images illustrating immunofluorescence staining of CD20, CD30 and LENG1 in non-lesional germinal center regions and tumor areas. Arrows indicate HRS cells. (J) Quantification of staining intensity of LENG1 in GC, HRS and SLL as (H). F (2, 40108) = 3054, **** P < 0.0001. (K) Expression of LENG1 determined by MS analysis (similar as in F). F (2, 9) = 217.7. (L) Representative immunohistochemistry images of LENG1 in human lymph node tissues, including both normal and lymphoma cases, sourced from the HPA database. (M) Kaplan-Meier analysis to pan-cancer in TCGA using UCSC Xena, with a focus on LENG1 (see main text). The p-values reflect comparisons between two groups through univariate analysis using the log-rank test.

In principal component analysis (PCA), the proteomes of the GC, HRS, and SLL populations clustered distinctly, implying substantial proteomic differences among the three populations. Specifically, the GC populations separated more from HRS than from SLL populations (**Figure 2C**). A total of 792 proteins were significantly differentially expressed between HRS and SLL (**Figure 2D**). In line with this, only 23 proteins changed commonly between HRS compared to GC and SLL compared to GC (**Figure 2G**). These findings suggest that, as expected, HRS and SLL are clonally unrelated entities, each exhibiting unique proteomic characteristics and expression profiles.

To further validate our single-cell type microdissection and proteome analyses, we selected three highly altered protein candidates for closer examination (**Figure 2A-B; 2D**). Interferon Regulatory Factor 4 (IRF4) emerged as the most differentially expressed protein in HRS compared to SLL (**Figure 2D; 2F**). This protein also served as a diagnostic marker of cHL ^25^ (**Figure 2G**), exhibiting consistent expression levels with immunofluorescence staining quantification (**Figure 2F; 2H**). This concordance between MS and immunofluorescence data reinforced the reliability of IRF4 quantification. SKI Like Proto-Oncogene (SKIL) was one of the top most differential proteins in SLL compared to HRS (**Figure 2D; Supplementary figure 3B**). Again, MS-based quantification matched closely with immunofluorescence staining quantification of SKIL (**Supplementary figure 3C-D**). Finally, Leukocyte Receptor Cluster Member 1 (LENG1) prominently stood out as the top highly differentially expressed protein in SLL compared to HRS, congruent with immunofluorescence staining (**Figure 2D, I-K**). The HPA database provided additional support by confirming elevated LENG1 protein levels in lymphoma compared to normal lymph nodes (**Figure 2L**).

We extended our investigation to assess the broader impact of LENG1 downregulation through a pan-cancer analysis using the University of California Santa Cruz (UCSC) Xena and TCGA dataset ^26^. This revealed that a decrease in LENG1 expression significantly correlated with worsened overall survival (**Figure 2M**). Together these validations provided robust evidence for the accuracy of our single-cell type microdissection and proteomics measurements. Furthermore, it identified potential prognostic markers like LENG1 for future investigations.

### Molecular alterations indicate potential targets and personalized treatments

The highly upregulated diagnostic markers, such as CD30 and IRF4 between HRS and GC, suggest potential targets for therapy (**Figure 1G**; **2G; 3A**). For instance, Lenalidomide, a thalidomide analogue currently under clinical trials for lymphoma according to the ClinicalTrials.gov dataset, exerts its pharmacological effects by modulating various cellular targets, including IRF4 (**Figure 3B**). Altogether, the differential protein expression analysis described above, revealed that compared to GC, HRS cells showed significant changes in 752 proteins, while SLL cells exhibited changes in 80 proteins (**Figure 3A; 3C**). This highlights the scope of potential opportunities for personalized targeted treatments for each of the tumors.

**Figure 3.**
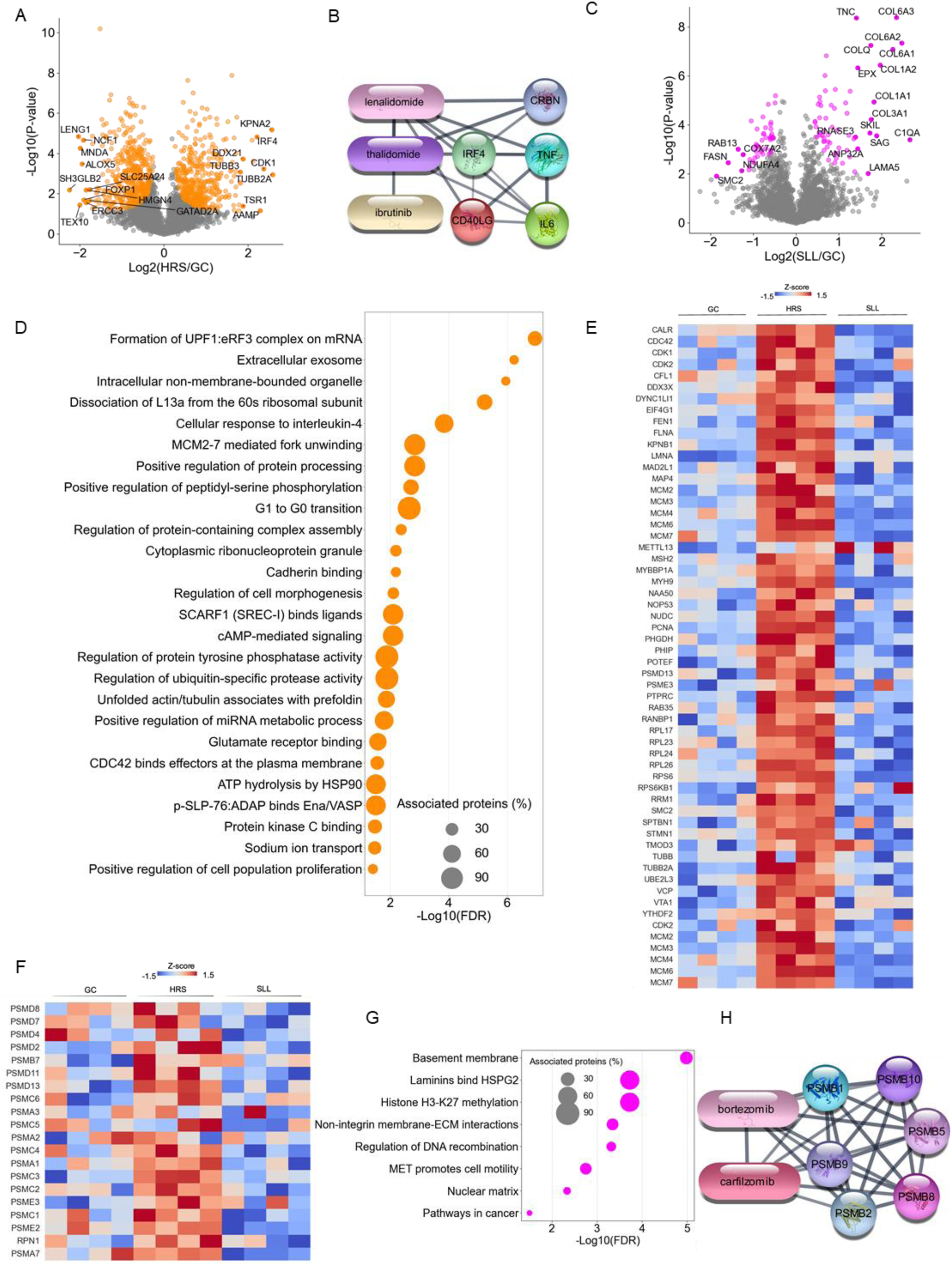
Identification of molecular changes, pathways and potentially druggable targets. (A) Volcano plot comparing the proteomes of HRS and GC. Orange dots represent significantly differentially expressed proteins, and proteins with largest fold-change are annotated with their names. Statistical significance was determined using two-sided *t*-test (FDR<0.05, S0=0.1). (B) Interaction between compounds and proteins using STITCH and STRING (see Methods). Interactions are visualized in Cytoscape, representing nodes with gray lines connecting them, where thicker lines signified stronger associations. IRF4 and related proteins appear as circles, while compounds appear as ovals in the visual representation. (C) Volcano plot comparing the proteomes of HRS and GC. Magenta dots represent significantly differentially expressed proteins as in (A). (D) Gene Ontology (GO) enrichment analysis on the upregulated pathways in HRS involving iterative comparisons and functional grouping of these pathways. The plot illustrates -log10 (FDR of term) values, with circle size denoting the percentage of associated upregulated entities in HRS compared to the background entities. (E) Heatmap illustrating entities participating in the cell cycle process pathway identified through GO enrichment analysis. GC, HRS, and SLL are represented with four replicates each, with Z-scores depicted using a gradient color scale. (F) Heatmap of the proteasome subunit proteins. (G) GO enrichment analysis of the upregulated pathways in SLL similar to (D). (H) Interaction between compounds and proteasome proteins.

Our MS-based analyses allowed us to identify specific cellular processes as potential targets for personalized medicine. For instance, we observed aberrations in cell cycle processes in HRS populations, including dysregulated activity of ubiquitin-specific protease/MCM (Mini-Chromosome Maintenance) proteins, dysregulation in the G1 to G0 transition, and increased cell proliferation. Furthermore, protein expression and processing was broadly dysregulated in HRS cells, which encompassed changes in protein tyrosine phosphatase/protein kinase C activity, miRNA metabolic processes, protein-containing complex assembly, ATP hydrolysis by HSP90, formation of UPF1:eRF3 complexes on mRNA, and the dissociation of L13a from the 60s ribosomal subunit. Additionally, we noted perturbations in cell movement within the HRS population, including disruptions in p-SLP-76:ADAP binding, CDC42 binding, and cell morphogenesis (**Figure 3D-E; Supplementary figure 4A-B**). Conversely, our analyses of the SLL population revealed aberrations in the regulation of lamin binding, H3K27 methylation, extracellular matrix (ECM) interaction, MET-promoted mobility, and DNA recombination. These findings suggest potential mechanisms underlying cancer development and progression specific to each of the tumors, indicating the potential therapeutic strategies targeting cell proliferation, protein turnover and processing, ECM interaction, and mobility.

The observed aberrations in cell cycle processes within the HRS population suggest potential sensitivity to chemotherapy (**Figure 3D-E**). The above-mentioned upregulated expression and activity of MCM proteins in HRS cells indicate the potential utility of MCM inhibitors (**Figure 3D-E**). Likewise, the upregulation of proteasome subunit proteins, including PSMA/B/C/D/E, in HRS cells (**Figure 3F**), points to proteasome inhibitors which are well established in the treatment of cHL. For instance, Bortezomib and Carfilzomib target PMSB proteins, and are currently in clinical trials for lymphoma treatment ^27^ (ClinicalTrials.gov) (**Figure 3G).** Moreover, the activation of ECM interaction, coupled with the expression of specific collagen in the SLL population, may make ECM-targeted therapies attractive ^28^ (**Figure 3H**). To investigate whether collagen originates exclusively from the ECM or if lymphoma cells also express collagen, we conducted Collagen I staining and observed it in SLL cells and not in GC cells (**Supplementary figure 3E**). For SSL cells, our findings point towards targeting H3K27 methylation and MET, a receptor tyrosine kinase (RTK) (**Figure 3H**).

The activated cellular response to interleukin (IL)-4 in HRS may reflect a mechanism of immune evasion, because activation of this pathway can suppress anti-tumor immune responses ^29^ (**Figure 3D**; **Supplementary figure 4C**). Additionally, inhibition of IL-4 signaling reportedly sensitizes HRS to chemotherapeutic drugs in a chemo-resistant cell line model ^30^, making this a promising approach for managing chemo-resistance.

In summary, a comprehensive treatment strategy could encompass chemotherapy and targeted therapies, including MCM inhibitors and proteasome inhibitors for cHL, as well as H3K27 methylation inhibitors and RTK inhibitors for SLL. Additionally, IL-4 inhibition may represent a valuable approach for addressing chemo-resistance in this case.

## Discussion

In this study, we utilized a composite lymphoma case to explore the potential of DVP for advancing precision oncology. Several key findings emerge from this case: 1) robust single-cell-type proteome analysis: our study demonstrates the accuracy and robustness of single-cell microdissection and MS evaluation; 2) distinct proteome profiles: we reveal that HRS and SLL populations exhibit distinct proteome profiles, indicative of divergent clonal development and progression mechanisms; 3) dysregulated pathways: in the HRS population, we observe disruptions in cell cycle processes, protein expression and processing, and cell mobility. Conversely, we note aberrations in lamin binding, H3K27 methylation, ECM interaction, MET-promoted mobility, and DNA recombination in SLL; 4) therapeutic targets: our analysis suggests promising therapeutic avenues encompassing chemotherapy and repurposed targeted therapies, including MCM inhibitors and proteasome inhibitors for cHL, along with H3K27 methylation inhibitors and receptor tyrosine kinase (RTK) inhibitors for SLL.

DVP facilitates the measurement of a patient’s disease-specific single-cell-type proteome from a single tissue slide, offering a wealth of information even from limited biopsy samples. With this technology, pathologists and clinicians can identify and evaluate predictive biomarkers and explore disease progression mechanisms, potentially leading to the discovery of new therapeutic targets. Notably, the validation of example proteins such as IRF4, LENG1, and SKIL in our study underscores the accuracy of our single-cell microdissection and MS evaluation procedures. Our *in silico* analysis suggests that LENG1 may positively impact overall survival, warranting further research into the roles of SKIL and LENG1 in composite lymphoma.

Proteomics indicated that the coexistence of cHL and SLL in this composite lymphoma case is unlikely to result from the transformation of SLL into cHL, which is also supported by the fact that *de novo* composite lymphoma with cHL in CLL/SLL patients is exceptionally rare. Instead, in general it is more plausible that SLL develop in HL patients or the development of HL in SLL patients may be related to exposure to immunosuppressive chemotherapy ^31, 32^. In our case, however, the patient had not received chemotherapy prior to the initial diagnosis. Richter transformation, in the form of the cHL variant, is typically suspected when a patient with SLL experiences a sudden onset of B symptoms. In contrast, here the clinical presentation of cHL indicated localized disease with a small tumor burden, supported by PET scan results, the absence of lymphoma growth, and the lack of B symptoms (**Methods**).

Our findings highlight promising therapeutic opportunities. Notably, IRF4 exhibited high expression in cHL, and Lenalidomide, a drug currently undergoing clinical trials for lymphoma treatment targets IRF4 amongst other proteins. The conclusive evidence of clonal unrelatedness helped guide the clinician’s decision to recommend chemotherapy. ABVD (Adriamycin, Bleomycin, Vinblastine, Dacarbazine) chemotherapy is widely recognized as a standard first-line treatment for cHL ^33, 34^. To date, the patient has undergone two series of ABVD treatment, resulting in complete remission of cHL as confirmed by PET-CT scans. Additionally, there has been a noticeable reduction in the size of other affected lymph nodes.

The activated pathways, such as the aberrant regulation of ubiquitin-specific protease activity, may indicate a potential mechanism through which cancer cells evade the normal process of protein degradation and turnover. This can lead to the accumulation of specific proteins that promote tumorigenesis or drug resistance. Furthermore, the activation of the IL-4 pathway suggests that targeting IL-4 could be beneficial in overcoming drug resistance. Our analysis has revealed not only the upregulation of the proteasome subunit protein PMSB but also PMSA/C/D/E in HRS cells. Therefore, it may be beneficial to consider the use of additional proteasome inhibitors for therapeutic intervention ^34^. In addition, proteasome inhibitors have demonstrated the ability to sensitize tumor cells to various anti-tumor therapies, indicating their potential role in managing chemo-resistance ^35^. JAK inhibitors could be valuable for blocking the downstream effects of IL-4 signaling since these drugs inhibit JAK kinases, which play a role in the IL-4 signaling pathway ^36^. Given the activation of IL-4 signaling in HRS, there may be potential benefits to using JAK inhibitors. In fact, studies have shown that the JAK inhibitor Ruxolitinib exhibits activity against HRS ^37^.

The identification of these targets holds promise for tailoring a personalized treatment for the patient. It also opens doors to potential enrollment in ongoing clinical trials and provides a means for monitoring treatment, especially regarding chemoresistance. In conclusion, this study offers valuable insights into the distinctive protein expression profiles of cHL and SLL within the context of composite lymphoma. The approach employed in this research can serve as a versatile framework applicable to similar cases, facilitating a deeper comprehension of the underlying biological mechanisms and the identification of potential targets for the development of novel treatments.

## Methods

### Case

A 71-year-old female patient with a medical history of diabetes and hypertension was admitted to Zealand University Hospital in Denmark with lymphadenopathy on the left side of her neck. The patient did not present with B symptoms or a previous cancer history. Histological examination of a left cervical lymph node revealed the presence of both cHL and SLL, as confirmed by two expert hematopathologists. The SLL cells were small and mature with clumped chromatin, and tested positive for CD20, CD23, CD5 and PAX5, while CD30, IMP3 and CD15 were negative and revealed a low proliferation index. On the other hand, the HRS cells were often binucleate, with large eosinophilic nucleoli and abundant cytoplasm and a pleomorphic cellular background including eosinophils and T-cells. The HRS cells stained positive for PAX5 (dim), CD30, IMP3, CD15, and P53, while CD20 was negative. Written consent to publish the case was obtained from the patient.

### Immunohistochemical staining on membrane slides

Membrane PEN slides 1.0 (415190-9041-000, Zeiss) were exposed to UV light for 1 hour and coated with Vectabond reagent (SP-1800-7, Vector Laboratories) according to the manufacturer’s protocol. FFPE lymph node tissue sections were cut at a thickness of 2.5 µm and mounted on treated PEN slides, left to dry at 37°C overnight, and heated at 60°C for 20 minutes to improve tissue adhesion. The sections were then treated to remove paraffin and hydrated by sequentially passing through xylene and decreasing concentrations of ethanol.

The Dako Omnis system was used for immunohistochemistry. Antigen retrieval was performed using Target Retrieval Solution Low pH (GV805, Agilent) and 10% glycerol (G7757, Sigma Aldrich) heated for 60 minutes at 90°C in a water bath. The tissue was then incubated with anti-CD30 clone CON6D/B5 (1:25, CM346C, Biocare Medical) for 30 minutes at 32°C. After washing and blocking of endogenous peroxidase activity, the Envision FLEX+High pH kit (GV800+GV821, Agilent) and Envision DAB+ Substrate Chromogen System (GV825, Agilent) were used for detection, following the manufacturer’s instructions. The use of DAB reaction products can create a shielding effect that helps prevent cross-reactivity with subsequent immuno-reagents ^38^. The second sequence involved incubation with anti-CD20 clone L26 (1:250, M0755, Agilent) using the same protocol, with reactions visualized using EnVision™ Flex Magenta Chromogen system (GV900, Agilent). Mayer’s hematoxylin was used for counterstaining.

For immunofluorescence, the sections were subjected to antigen retrieval in EDTA buffer (E1161, Sigma-Aldrich) with 10% glycerol at 90°C for 45 minutes. The tissue was then blocked with 5% BSA for 20 minutes at room temperature and incubated overnight at 4°C with primary antibodies, including anti-CD30 clone Ber-H2 (1:50, M0751, Agilent) with anti-SKIL antibody (1:200, HPA008472, Sigma-Aldrich) / anti-IRF4 antibody (1:100, 11247-2-AP, Proteintech) / anti-LENG1 antibody (1:500, HPA043541, Sigma-Aldrich) / Alexa Fluor® 488 Collagen I (1:1000, ab275996, Abcam) / CD3 (1:500, A0452, Agilent). Unconjugated primary antibodies were incubated with the corresponding Alexa Fluor®-labeled secondary antibodies at a 1:1000 dilution for one hour at room temperature. The sections were then incubated with eFluor™ 660 anti-CD20 clone L26 (1:50, 50-0202-80, Invitrogen). The slides were counterstained with DAPI and mounted with Thermo Fisher Diamond Antifade mounting media, then examined using an AxioScan7 microscope (Zeiss).

### Image analysis

Annotated training data based on CD20 and CD30 staining was supervised by an expert hematopathologist and was used to train a project-specific Mask R-CNN segmentation model ^3^. This model was integrated into BIAS software and then used to accurately segmented target compartments ^3^. The BIAS software was used to classify normal B cells from residual germinal center region, as well as HRS and SLL cells based on CD20 and CD30 staining (HRS cells are CD20^-^CD30^+^ while SLL/normal B cells are CD20^+^CD30^-^) and cellular morphological features.

To calculate the distance between HRS and SLL, we used the Pythagorean Theorem based on the x and y-coordinates of individual cells. For example, the distance between a test cell (x_test, y_test) and a given cell (x_given, y_given) can be calculated using the following equation:

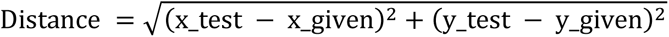

### LMD and peptide preparation

Using the Leica LMD7 system and an HC PL FLUOTAR L ×63/0.70 CORR XT objective, the cellular contours were precisely excised from membrane slides ^3^. The Leica Laser Microdissection V 8008 software was used for automated excision and collection of the contours. A surface area of 75000 μm^2^ of each sample was collected (around 500 HRS cells). To each sample well, 30 µL of 100% acetonitrile was added. The plate was sealed with sealing tape and vortexed for 10 seconds. The plates were then centrifuged at 2000 g for 3 minutes and vacuum dried at 60 °C. 4 µL of 60mM triethylammonium bicarbonate (TEAB) was added to each sample and heated at 95 °C for 1 hour in a thermal cycler (S1000 with 384-well reaction module, Bio-Rad) set to a constant lid temperature of 110 °C. 1 µL of 60% acetonitrile was added and the samples were heated at 75 °C for 1 hour. The samples were cooled and 4 ng of LysC (Wako) was added and incubated at 37 °C for 3 hours in the thermal cycler. 6 ng of trypsin (Sigma-Aldrich) was added and the samples were incubated overnight at 37 °C. The digestion was stopped by adding 1% v/v trifluoroacetic acid (TFA), and the samples were vacuum dried at 60 °C. 4 µL of MS loading buffer (3% acetonitrile in 0.2% TFA) was added, the plate was vortexed for 10 seconds, and centrifuged at 2000 g for 5 minutes. The samples were stored at -20 °C until they were ready for MS analysis.

### LC-MS/MS analysis

To generate a project-specific library for data-independent mass spectrometry (MS) analysis, we employed high-pH reversed-phase fractionation. The process involved fractionating peptides at pH 10 using the automated Opentrons platform for fraction collection, in conjunction with a nanoflow HPLC (EASY-nLC 1000 system, Thermo Fisher Scientific). Subsequently, we separated 20 µg of purified peptides obtained from bulk lymph node tissue. This separation occurred on an analytical column (250 µm x 30 cm, 1.9 µm, PepSep™, Bruker Daltonics) through a 100-minute gradient, with an exit-valve switch taking place every 30 seconds. Ultimately, we concatenated the eluted peptides into 48 fractions. The peptide fractions were vacuum dried and reconstituted in MS loading buffer. MS analysis was performed using an EASYnLC-1200 system (Thermo Fisher Scientific) interfaced to a modified trapped ion mobility spectrometry quadrupole time-of-flight mass spectrometer (timsTOF Pro, Bruker Daltonik) with a nano-electrospray ion source (CaptiveSpray, Bruker Daltonik). The peptides were loaded onto a 50 cm in-house-packed HPLC column (75-µm inner diameter packed with 1.9-µm ReproSil-Pur C18-AQ silica beads) and separated using a linear gradient from 5-30% buffer B (0.1% formic acid and 80% ACN in LC-MS-grade water) over 55 minutes, followed by an increase to 60% for 5 minutes and a 10-minute wash in 95% buffer B at a flow rate of 300 nL/min. Buffer A consisted of 0.1% formic acid in LC-MS-grade water. The column temperature was kept constant at 60 °C using an in-house-made column oven. The total gradient length was 70 minutes. MS analysis was performed in either data-dependent (ddaPASEF) or data-independent (diaPASEF) mode as described by Mund and Brunner et al. ^3, 39^.

### Data analyses

The raw files of the project-specific spectral library acquired in ddaPASEF mode were processed using MSFragger (version 18.0)^40^, with the exception that cysteine carbamidomethylation was not included as a fixed modification. The resulting library file was used in DIA-NN (version 1.8.1) ^41^ to analyze the raw files of diaPASEF measurements. The samples in diaPASEF mode were subjected to analysis using DIA-NN, which employed a library-based approach against the UniProt database, including isoforms (2019, UP000005640_9606). The settings comprised trypsin specificity with allowance for one missed cleavage, a precursor FDR threshold of 1%, accuracy of 15 ppm, and activation of the ’match between runs’ feature. N-terminal methionine excision, methionine oxidation, and N-terminal acetylation were selected, while a maximum of 2 variable modifications were permitted. The proteome datasets underwent filtration to retain samples with at least 70% valid values (proteins exhibiting more than 30% missing values were excluded from subsequent statistical analyses). Any remaining missing values were imputed using the K-Nearest Neighbors (KNN) imputation method ^42, 43^. The intensity values for all samples underwent a logarithmic transformation (log2) and were subsequently utilized for principal component analysis (PCA) and the identification of differentially expressed proteins. Visualization of these proteins was achieved through volcano plots and heat maps, using Python and R for analysis. For multi-sample comparisons (ANOVA) or pairwise proteomic analyses (two-sided unpaired t-test), a protein was considered significantly differentially abundant if the FDR-adjusted P-value was less than 0.05.

The Kaplan-Meier method was used to estimate overall survival, and differences were assessed using the log-rank test. The Search Tool for Interactions of Chemicals (STITCH) database was used to retrieve the bioactive components and targets of chemicals being studied in clinical trials for lymphoma, as listed in the ClinicalTrials.gov dataset ^44^. Protein-protein interactions were analyzed using the Search Tool for the Retrieval of Interacting Genes/Proteins (STRING) database ^45^, and the Cytoscape software (version 3.9.1) was used to create plots of compound-target interactions ^46^. Ontology enrichment analysis was conducted on the liver proteomics dataset using ClueGo (v2.5.10), a Cytoscape (v.3.1.1) plug-in, with default settings. A customized reference set comprising 3257 unique genes quantified in this study was employed in Fisher’s exact test. Term significance was corrected using the Benjamini–Hochberg method with an FDR threshold of less than 1%. This analysis included the activation of both Gene Ontology term fusion and grouping. Significantly enriched terms and the associated proteins are detailed in the Supplementary section.

## Acknowledgements

We gratefully acknowledge the contributions of Maria Wahle, Elena Krismer, Dr. Constantin Ammar, Dr. Lylia Drici, Dr. Juanjuan Wang and Dr. Lili Niu, who provided helpful discussions on the mass spectrometric analysis and data visualization. We also thank Dorthe Strue-Nielsen and Linda Bagger Hansen for technical support. We are grateful to the patient who generously allowed us to share the medical history in this case study and helped us to gain valuable insights into this rare condition. This work was supported by the Novo Nordisk Foundation (grant agreements NNF14CC0001 and NNF15CC0001).

## Contributions

Conceptualization: M.M, X.Z., L.M.R., A.M.; Methodology: X.Z., A.M., M.B.; Experiments: X.Z., M.B.; Data analysis: X.Z.; Supervision: M.M; Funding acquisition: M.M; Writing original draft: X.Z. All the authors reviewed and edited the manuscript.

---

**Supplementary figure 1.**
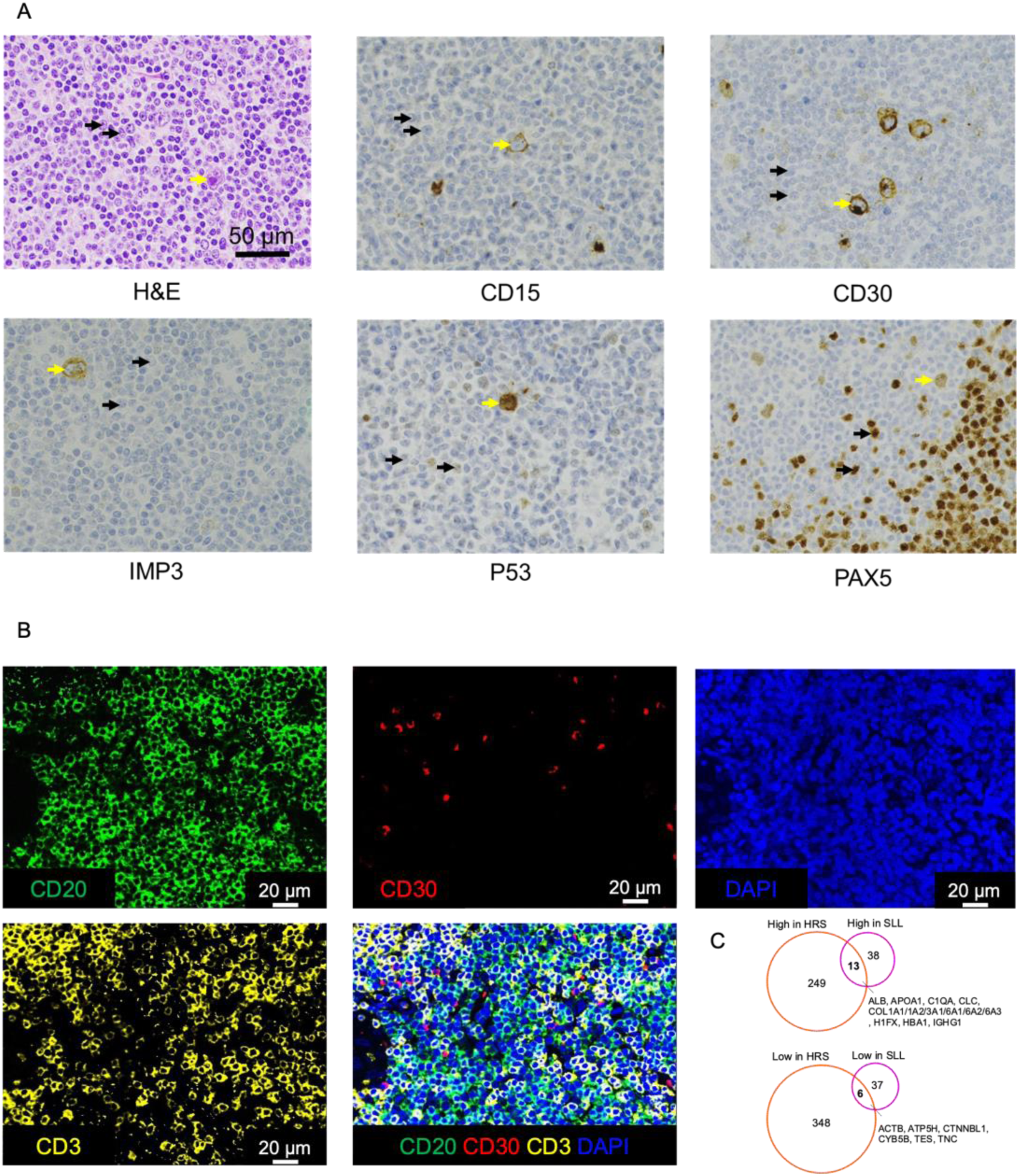
A case with composite small lymphocytic lymphoma (SLL) and classical Hodgkin lymphoma (cHL). (A) Representative images showcasing the immunohistochemistry staining of diagnostic markers of the lymphoma case. The yellow arrow highlights examples of HRS cells, while the black arrow points to examples of SLL cells. (B) Representative images illustrate immunofluorescence staining of CD20, CD30 and CD3 in tumor areas. (C) Venn diagram analysis of proteins significantly differentially expressed in HRS or SLL compared to GC. 6 overlapping proteins that are expressed at lower levels in both HRS and SLL compared to GC and 13 overlapping proteins that are highly expressed in both HRS and SLL compared to GC are shown. Statistical significance was determined using two-sided *t*-test (FDR<0.05).

**Supplementary figure 2.**
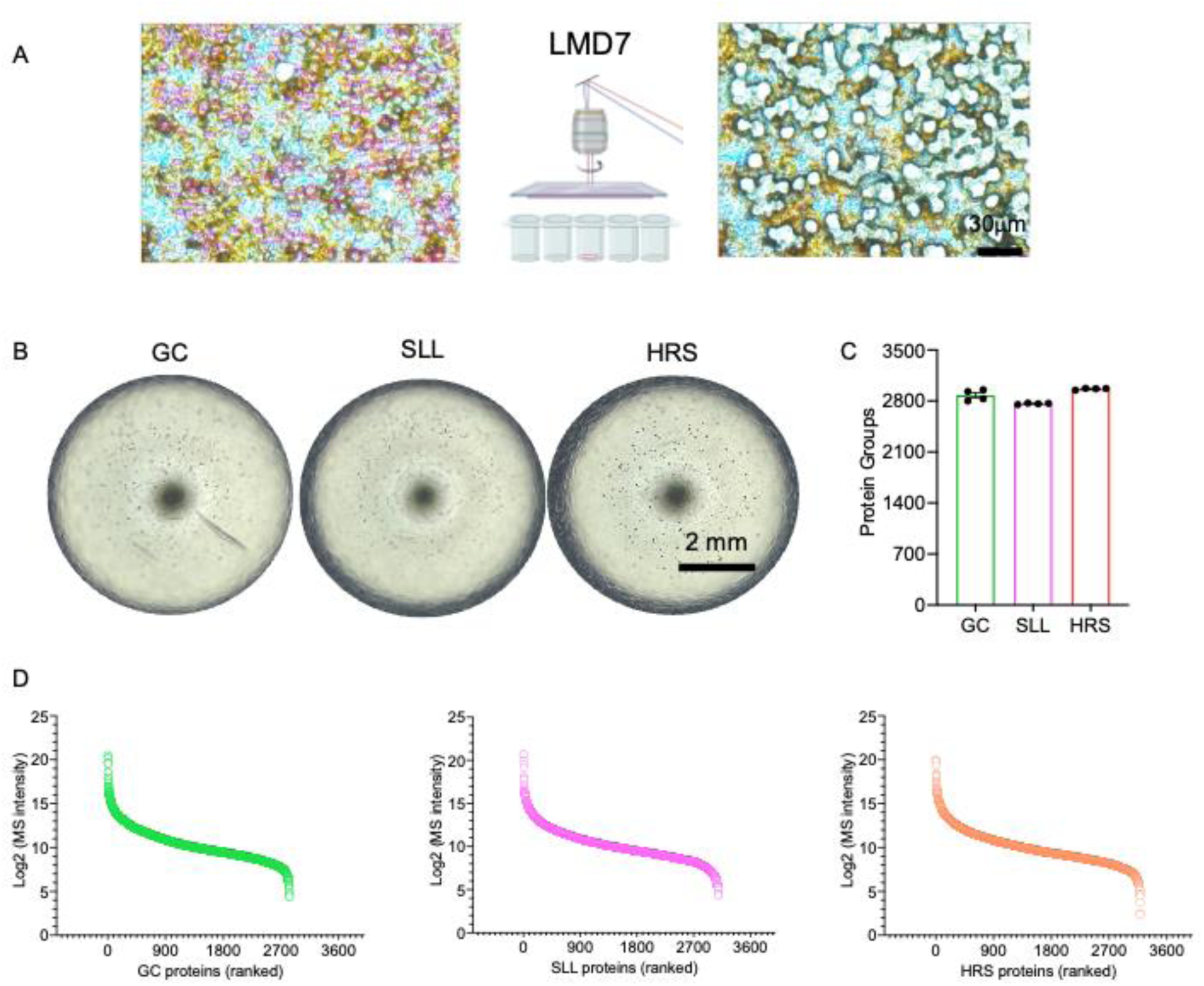
Laser microdissection and mass spectrometry analysis of GC, HRS and SLL populations. (A) Single cells of each GC, HL and SLL populations were isolated using automated laser microdissection and collected into 384-well plates. The panel displays cell contours before and after microdissection. (B) Three representative images of a well containing the collected cells are shown. (C) The number of proteins identified in GC, HRS and SLL samples. (D) Protein ranking plot of proteins identified in GC, HRS and SLL samples. Proteins are ranked by transformed intensity values detected by MS, which directly represent the protein abundance.

**Supplementary figure 3.**
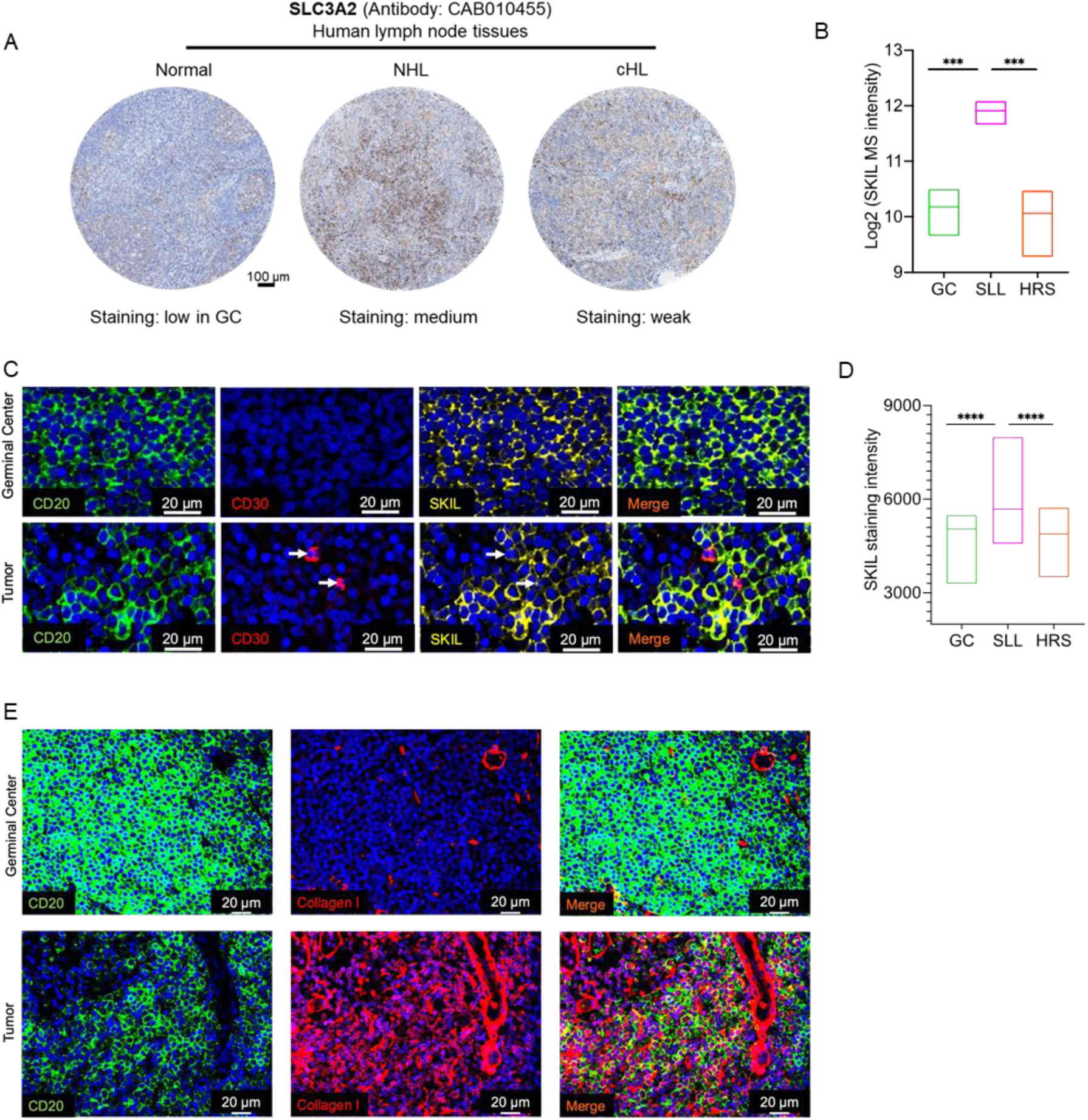
Validation of example proteins identified by MS. (A) Representative immunohistochemistry images of SLC3A2 in human lymph node tissues, including both normal and lymphoma cases, sourced from the HPA database. (B) Expression of SKIL, as determined by MS analysis. The box plot displays the distribution of the data from the minimum to the maximum intensity. The horizontal line within each box represents the mean value of the intensity. Statistical significance was determined using one-way ANOVA. F (2, 9) = 20.04, **** P < 0.0001. (C) Representative images illustrate immunofluorescence staining of CD20, CD30 and SKIL in non-lesional germinal center regions and tumor areas. Arrows indicate HRS cells. (D) Quantification of staining intensity of SKIL in GC, HRS and SLL. The box plots display the distribution of the data from the minimum to the maximum intensity. The horizontal line within each box represents the median value of the intensity. Statistical significance was determined using one-way ANOVA. F (2, 41965) = 4583, *** P < 0.001. (E) Representative images illustrate immunofluorescence staining of CD20 and Collagen I in tumor areas.

**Supplementary figure 4.**
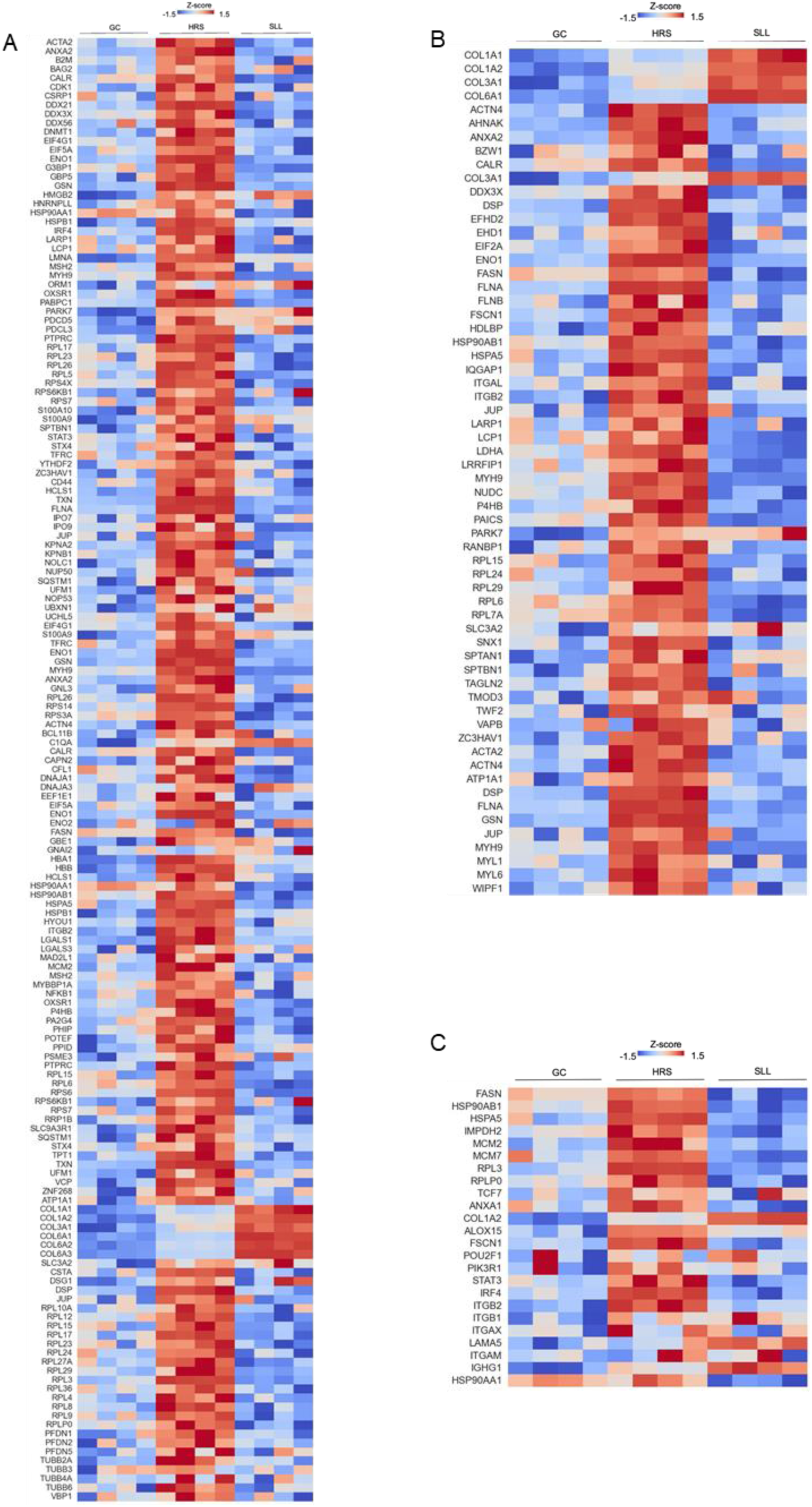
Identification of molecular changes and evaluation of potential druggable targets. The heatmaps display the entities involved in the protein processing and regulation (A), cell mobility (B), and IL-4 (C)-related signaling pathways, showing three groups of GC, HRS, and SLL, each with four replicates, and Z-scores are depicted using a gradient color scale.

